# Biological warfare between two bacterial viruses in a genomic archipelago sheds light on the spread of CRISPR-Cas systems

**DOI:** 10.1101/2023.09.20.558655

**Authors:** Alejandro Rubio, Andrés Garzón, Antonio Moreno-Rodriguez, Antonio J. Pérez-Pulido

## Abstract

CRISPR-Cas systems are acquired immunity systems of bacteria and archaea that prevent infection by phages and other mobile genetic elements. It is currently known that this defense system has also been co-opted by viruses. These viruses could use CRISPR-Cas systems to compete against other rival viruses. We have discovered a virus in the bacterium *Acinetobacter baumannii* that incorporates a CRISPR-Cas system into an integration hotspot of the host genome. Once integrated, this could prevent the infection of the most frequent viruses in this bacterial species, especially one that competes with the CRISPR-Cas system itself for the same integration site. This latter virus is prevalent in strains of the species belonging to the so-called Global Clone 2, which causes the most frequent outbreaks worldwide. Knowledge of this new viral warfare using CRISPR-Cas systems, known to limit the entry of antibiotic resistance genes into bacteria, could be useful in the fight against the infections they cause. But it would also shed light on the way in which these defense systems expand in bacteria.

## Introduction

*Acinetobacter baumannii* is a bacterium that causes endemic infections and hospital-acquired outbreaks with a high mortality and morbidity rate. For this reason, the World Health Organization recently classified it as “priority 1” on a list of pathogenic bacteria for which the development of new antibiotics is urgent (1). The bacterium can colonize the human body and even resist in the hospital environment, and opportunistically causes pneumonia, urinary tract and wound infections and, occasionally, bacteremia (2, 3). Much of the success of *A. baumannii* can be attributed to its genome plasticity, which mutates rapidly under adverse conditions and stress (4).

Strains of *A. baumannii* can be classified according to clonal groups based on several conserved genes (5), and two adjacent clonal groups reflect genomes arising from a recent common ancestor. In one of the reference schemes (the Institut Pasteur MLST), ST2 corresponds to Global Clone 2, which is by far the dominant group of this species, causing a large proportion of outbreaks worldwide (6).

As found in 40% of bacterial genomes, *A. baumannii* possesses the acquired CRISPR-Cas defense system against bacterial viruses (bacteriophages, known as phages) and other mobile genetic elements (7). However, this system appears in only 13% of its sequenced genomes (8, 9). These CRISPR-Cas systems are mainly targeted against phages, but also against plasmids to a lesser extent (10). Finally, it has been discussed that they could also be targeted against phage-plasmids (8), which are phages that can act as plasmids under certain conditions (11).

CRISPR-Cas systems are highly diverse prokaryotic acquired immunity systems, currently classified into 2 classes, with multiple types and subtypes, which differ in the architecture of their components and their sequence specificity (12). They are composed of genes that are part of the different stages of this immune system and are generically called *cas* (CRISPR-associated genes) (13). When the phage DNA enters the host, its sequence is cleaved into fragments that are added to the CRISPR array by the Cas1 and Cas2 proteins in a process called adaptation. During this step, phage sequences of about 32 nucleotides, known as spacers, are recognized and stored in a CRISPR array, separated by sequence repeats of similar length. The CRISPR array is then transcribed during the expression stage, resulting in transcripts that are cleaved into their corresponding spacers. Finally, during the interference stage, the spacer recognizes the complementary sequence in the target phage, called protospacer, which leads to the elimination of the viral infection by cleaving the phage DNA. This last step requires a nuclease, which in class I CRISPR-Cas systems is a protein complex in which the Cas3 protein stands out as the nuclease responsible for cleaving the phage genome. In addition, for the identification of the protospacer, it is essential to recognize a sequence of a few nucleotides (often only 2 or 3) just at the 5’ position of the protospacer, known as protospacer adjacent motif (PAM) sequence (14).

*A. baumannii* mainly presents class I CRISPR-Cas systems, divided into 2 type I subtypes: I-Fa (also classified as I-F2) and I-Fb (also classified as I-F1) (8, 15–17). Remarkably, Global Clone 2 does not usually present CRISPR-Cas systems (9).

Phage-encoded CRISPR-Cas systems have also been discovered in recent years (18). The most characterized of these is the phage ICP1 of *Vibrio cholerae*, which recognizes another phage with which it competes (known as satellite phage). This satellite phage needs ICP1 to complete its lytic cycle, and ICP1 uses an I-F type CRISPR-Cas system to defend against the satellite phage (19).

Phages that escape the bacterial defense systems and integrate into the bacterial genome, entering the so-called lysogenic cycle, can remain as prophages until they resume lytic cycle, in which they replicate and can leave the bacterium (20). These prophages are usually integrated into tRNA and tmRNA genes, as well as other genomic elements including integrative plasmids, and genomics islands. It is facilitated by the specificity of their integrases (21). Genomic islands are sometimes grouped in so-called archipelagos, which have been little studied so far (22, 23).

We have found an archipelago in an *A. baumannii* tmRNA gene, prominently featuring a prophage that appears to compete for that site with one of the bacterial CRISPR-Cas systems. Remarkably, this same CRISPR-Cas system is sometimes carried by a phage-plasmid. All of this suggests that both phages may be competing for the bacterium, and that the phage-plasmid may leave the CRISPR-Cas system integrated into the bacterium to gain the advantage in that armed race between phages.

## Materials and methods

### Pangenome construction

The pangenome used was the one constructed with 9,696 genomes in Rubio et al., 2023 (8). All genomes are structurally and functionally annotated, as well as the reference genes of the pangenome itself. The CRISPR-Cas systems of each genome, as well as the spacers and repeats of the CRISPR arrays were also obtained from the aforementioned work. In some cases the CRISPR-Cas systems were checked manually, especially in the I-Fa subtype, finding some additional ones that were mostly incomplete (lacked CRISPR array or some or most of the cas genes). The latter systems were labeled as ambiguous or incomplete.

### Phage search

Prophages were searched in all the genomes using Phigaro version 2.3.0 with default parameters and the *abs* mode (56). Only phages ≥9Kb in length were considered. Then, prophages were grouped by similarity using the MeShClust version 3.0 program with the parameters *-v* (total initial sequences) and *-b* (1/4 of total initial sequences). To combine similar phage genomes with the sequence oriented in the two possible reading directions, we performed a similarity search with BLASTN by comparing the reference sequences of each cluster with each other. Then, clusters whose reference sequence shared at least 90% identity and coverage (with respect to the shortest sequence) were combined. Finally, the shortest sequence from each final group was selected as the reference sequence.

The phage-plasmid was searched for in contigs of no more than 130,000 Kb, in which it was found with at least 90% identity and 70% sequence coverage with respect to a reference sequence that had it (GenBank: NZ_UFOI01000014.1), to which the CRISPR-Cas system was masked. CRISPR-Cas I-Fa and restriction-methylation systems were manually checked on these sequences, searching for genes *hsdR*, *hsdS* and *hsdM*.

The full version of the prophage DgiS1 was obtained by taking the nucleotide sequence from the phage integrase (gene homologs of GenBank: CU468230.2) to the direct repeat of the *ssrA* gene (GTTCAACTCCCGCCATCTCCACCA). Incomplete versions of the prophage DgiS1 were taken by searching for the phage integrase and checking that the genome had at least 40% of the complete phage sequence.

The rest of the genomic islands integrated in *ssrA* were obtained by searching for the core gene most likely to be the one that should appear next to *ssrA*, and extracting the different gene clusters that could be found in between them.

### Phage annotation

The phage-plasmid was found in independent contigs in the genomes analyzed. After performing a similarity search on the NCBI-Blast web portal, several plasmids were found with more than 90% identity and coverage, with and without the CRISPR-Cas I-Fa system. PHASTEST was used to annotate the genome of this phage (24).

VPF-Class tool (no version) was used to annotate the putative genre of the prophage integrated in *ssrA* (P1virus-like) (25). PhANNs was used to annotate the putative viral genes of the different genomic islands integrated in *ssrA* (26). The gene class assigned to each viral gene was assigned as the one with the highest score.

### Molecular phylogenies

Molecular phylogenies were performed using the assembly of complete bacterial or phage genomes, always using an outgroup to root the trees. Phylogenies were performed with the web tool Realphy 1.13 (27) and the trees were visualized with the ggtree library of the R programming language.

Due to the high sequence similarity of the phage-plasmids carrying the CRISPR-Cas I-Fa system, the processing of their sequences was homogenized. The raw data from these sequences were downloaded using SRA Toolkit v3.0.8 (https://github.com/ncbi/sra-tools). Next, each genome was reassembled using Unicycler v2022 (28). Finally, the contigs containing the single phage-plasmid contigs were selected by searching for the *unknown5757* gene in the newly assembled genome. The average length of these contigs was 110 kb.

### Retrieval of CRISPR repeats and spacers

Initially, the CRISPR array repeats were extracted from the output of the CRISPRCasFinder tool (29). Their orientation was taken from the ’potential_direction’ output attribute. Since most of the repeats had a value equal to ’unknown’, the different variants of the repeats were reviewed manually and those that were similar were grouped together, transforming those that were in reverse-complementary direction. In this way, the direction of all CRISPR arrays could be determined.

Subsequently, the spacers were extracted, already with the direction suggested by the flanking repeats. In this way, sequence logos could be made using the ggseqlogo library of the R language.

### Discovery of protospacers

Protospacers were searched by performing a similarity search with BLASTN and the *blastn-short* option turned on, using a threshold of ≥95% sequence identity and 100% spacer coverage.

## Results

### *Acinetobacter baumannii* has two different subtypes of CRISPR-Cas I-F systems

We analyzed an *A. baumannii* pangenome composed of 9,696 genomes, in which we searched for CRISPR-Cas systems. Two different configurations of CRISPR-Cas systems were found, both type I-F. Specifically, 9.5% of the genomes had the I-F1 subtype (also referenced as I-Fb in this species), 3.5% presented a system similar to the I-F2 subtype (also referenced as I-Fa), and 1.6% were classified as ambiguous, since they only had isolated CRISPR arrays or some isolated *cas* genes (Fig. 1A). The I-Fa system presented 2 independent CRISPR arrays, as previously known (9). The difference between both subtypes was given precisely by the number of CRISPR arrays, but also by the order of the *cas* genes and the genomic integration site. Furthermore, the amino acid sequence of the encoded Cas proteins only shares about 30% identity between the homologs of both subtypes. In addition, the Cas2_3, Cas8f and Cas7f proteins of the I-Fa subtype are notably shorter than the corresponding homologous proteins of the I-Fb system.

**Fig. 1.**
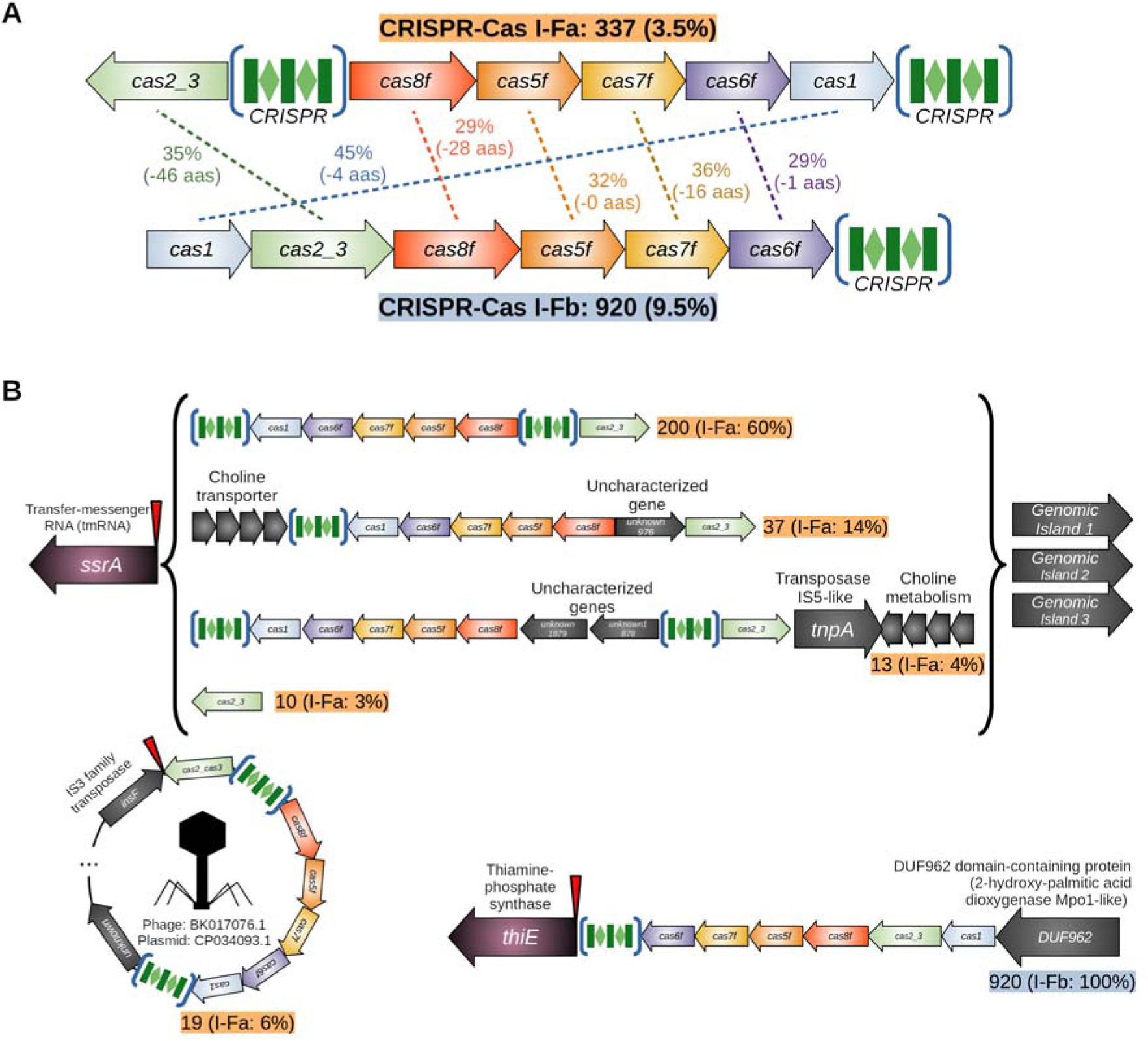
CRISPR-Cas I-F systems in *Acinetobacter baumannii*. (A) Gene structure of the I-Fa and I-Fb CRISPR-Cas systems. Next to the type of CRISPR-Cas system, the number of genomes that have it and the percentage with respect to the total number of genomes analyzed are shown. In the middle of both CRISPR-Cas systems, the percentage of identity between the homologous proteins of both systems is shown, and in parentheses the number of amino acids that the Cas proteins of subtype I-Fa had fewer than I-Fb. (B) Genomic regions in which CRISPR-Cas systems are found. The I-Fa subtype is mostly integrated behind a tmRNA called *ssrA*. But the latter is also found downstream of *ssrA*, under a cluster of genes including a choline transporter, together with a transposase and two uncharacterized genes, with only one of the *cas* genes (*cas2_3*). Finally, it also appears in a circular molecule, together with another transposase. The latter configuration is also found in a plasmid in the NCBI RefSeq database (CP034093.1) and in several phages (BK017076.1; 30% coverage). The I-Fb subtype is always located between the *thiE* gene and an uncharacterized gene encoding a protein with the DUF962 domain. The red inverted triangles indicate the genomic islands insertion site.

The I-Fb system appeared always integrated at exactly the same site, between the *thiE* gene (encoding a Thiamine-phosphate synthase) and a gene encoding an uncharacterized protein (Fig. 1B). However, the I-Fa system is found to be integrated with different frequencies at different sites. In almost two-thirds of the genomes with this subtype, the CRISPR-Cas system is integrated near the 5’ end of the *ssrA* gene, a transfer-messenger RNA (tmRNA). In 14% of the genomes, the CRISPR-Cas I-Fa system is displaced by a cluster of 14 genes including a choline transporter. Less frequently, the system is interrupted by 2 uncharacterized genes and at the opposite end of *ssrA* is an IS5-like transposase, followed by the choline transporter cluster. Moreover, we found 10 genomes in which only the *cas2_3* gene appears next to the *ssrA* gene. In all these cases, other gene clusters are usually found downstream of the CRISPR-Cas I-Fa system, in the form of genomic islands, in addition to the one containing the choline transporter.

Finally, we found that the CRISPR-Cas I-Fa system also appeared in episomes of about 110 Kb, not assembled with the bacterial chromosome sequence, together with an IS3 family transposase and a number of other genes. When we searched for sequences similar to these episomes in the NCBI RefSeq database, we found several plasmids, which did or did not contain the CRISPR-Cas system (Suppl. Fig. S1). Furthermore, when we analyzed its sequence, we found that it could also be a phage (105 of its 112 genes are annotated as putative phage genes). This latter, together with the previous result, suggests that it could actually be a phage-plasmid that occasionally carries a CRISPR-Cas I-Fa system. We named this phage-plasmid as Phage-Plasmid Terminator of Other Phages (PPTOP), based on the defense system it carries.

The rest of the CRISPR-Cas I-Fa systems appear in other less frequent locations or their position cannot be determined, due to their fragmentation in the genomes analyzed. It is important to note that the *cas2_3* gene is the only one that appears in all the CRISPR-Cas I-Fa system configurations found. The protein it encodes has about 30% identity with that of the *Vibrio cholera* phage ICP1 (RefSeq:AGG09390.1), which was one of the first phages known to carry a CRISPR-Cas I-F system, and the best characterized to date (19, 30).

### The *ssrA* gene provides support for an archipelago for the integration of multiple genomic islands

Since tRNA and tmRNA genes are often hotspots for integration of genomic islands in bacteria, we wanted to know whether other elements, in addition to the CRISPR-Cas I-Fa system, were integrated into *ssrA*. Thus, when analyzing the genes that appear downstream of the *ssrA* gene, we found that other clusters of genes appeared integrated into it, displaying different frequencies (Fig. 2). The most frequent were two, the cluster containing the choline transporter, and another with a putative prophage. This prophage has a length of about 82 Kb, with a CG content of 36.5% (*A. baumannii* presents an average of 38.5%), has 90 genes, and is classified as a putative P1virus-like phage. A proof of the virus integration is that a direct repeat is always found at the end of it, with the same sequence as the 3’ end of the *ssrA* gene, probably representing a cognate att (phage attachment) sequence (see Materials and Methods). This prophage includes a toxin-antitoxin system that could help its persistence in bacteria, with a RelE/StbE family addiction module toxin and the corresponding mRNA interferase RelE/StbE antitoxin. The prophage is found in 4,597 genomes (47% of all genomes analyzed), although only about half of them were complete between the *ssrA* gene and the distal att sequence. Due to its high frequency, we named this prophage as Genomic Dominant Island in SsrA 1 (Prophage Acinetobacter DgiS1).

**Fig. 2.**
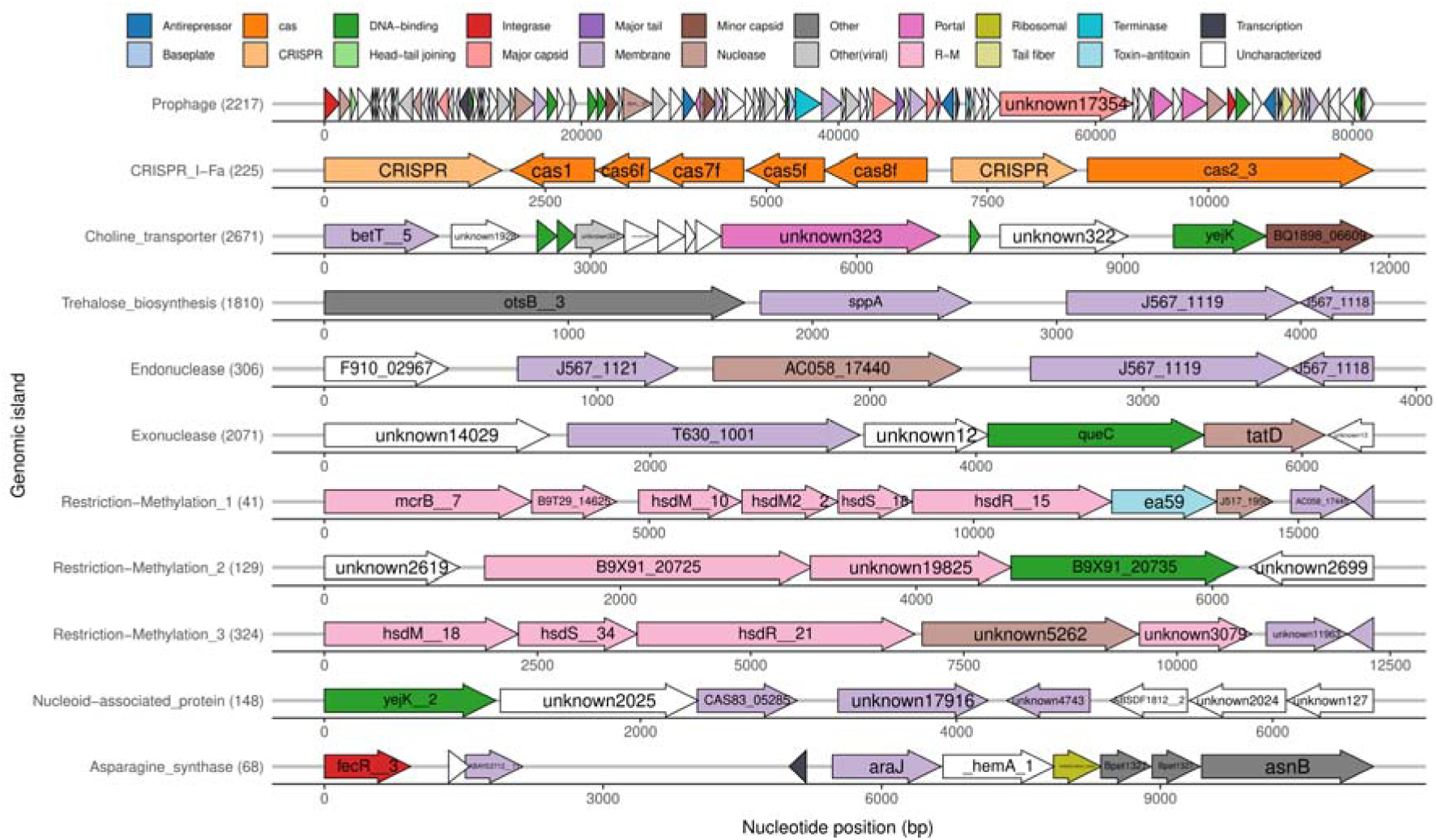
*ssrA* integrated genomic islands. Colors highlight annotated functions for genes in each genomic island. The different elements of the CRISPR-Cas I-Fa system are shown in orange. R-M corresponds to functions related to restriction-methylation systems. Some of the annotations correspond to viral functions: Baseplate, Head-tail joining, Integrase, Major capsid, Major tail, Minor capsid, Portal, Tail fiber, Terminase and Other(viral). The numbers in parentheses accompanying genomic island names represent the absolute frequency of that genomic island next to the *ssrA* gene.

Other genomic islands integrated at the same site are a cluster of 4 genes with trehalose biosynthesis genes, another with arginine synthesis genes, 2 especially frequent with endonucleases or exonucleases, and 3 different islands that seem to contain restriction-methylation systems, and that could be part of the bacterial defensome, as well as the CRISPR-Cas system. Some of the genes in these genomic islands are similar to phage genes, especially in the case of the DgiS1 prophage, in which several of them are predicted to be associated with the viral capsid.

### The prophage DgiS1 and the CRISPR-Cas system do not usually coexist in the same genome

Genomic islands of *ssrA* often appear displaced by others and may take up different positions between the *ssrA* gene and the core gene in *A. baumannii* (gene present in all the genomes of the species) that encodes a hypothetical lipoprotein (Fig. 3A). In fact, this bacterial core gene only appears next to *ssrA* in 3 of the genomes analyzed. Remarkably, the prophage DgiS1 appears to take up only the first position (next to *ssrA*), probably driven by the presence of the att site in the *ssrA* gene. On the contrary, the genomic island with the choline transporter may take up either the first or the second position. In fact, the prophage DgiS1 appears as the only genomic island in more than 2,000 genomes, or followed by other genomic islands in lower frequency (Fig. 3B). In contrast, the CRISPR-Cas system also appears alone or preceded by the island with the choline transporter. Notably, the prophage DgiS1 and the CRISPR-Cas I-Fa system do not usually appear together in the same genome, and when they do, the prophage appears mostly in its incomplete version (Fig. 3C). Specifically, they co-occur in 37 genomes (Fisher exact test, *p-value* = 7.39e-60), and only in one of them the CRISPR-Cas system appears next to the *ssrA* gene. It is also noteworthy that 31 of the genomes have both CRISPR-Cas I-Fa and I-Fb systems, almost the number that would be expected by chance (Fisher exact test, *p-value* = 0.92).

**Fig. 3.**
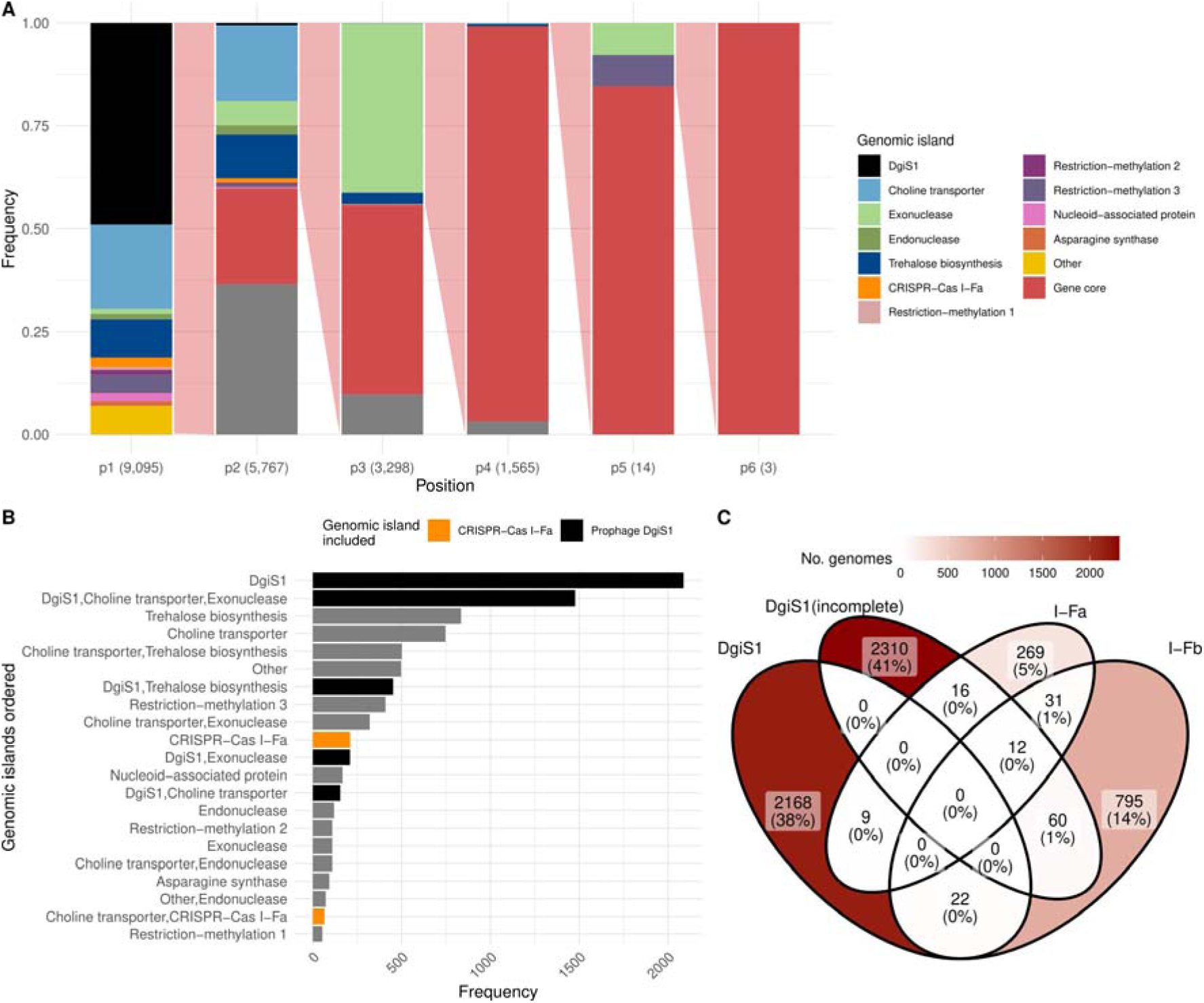
Position of the genomic islands integrated in the *ssrA* gene. (A) Relative frequency of each genomic island at each sequential position (p1 corresponds to the closest to *ssrA* and p6 to the farthest). ’Other’ indicates that the genomic island is none of those studied here. The core gene is the bacterial gene encoding a putative lipoprotein that should appear next to *ssrA* (gene homologs of GenBank: CAM88546.1), if there were no integrated genomic islands. When the genomic island could not be determined, it is highlighted in gray. Numbers in parentheses show the number of total genomic islands found at each position. Red shading between bars indicates the proportion of genomes with genomic islands at a given position that have additional genomic islands at the next position. (B) Frequency of the different combinations of genomic islands found. Only the most frequent combinations are shown. (C) Number of genomes having the DgiS1 prophage, or the CRISPR-Cas I-Fa or I-Fb systems, including group overlapping.

The non-coexistence of these two genomic islands could be due to the evolutionary divergence of the genomes containing them. To discard this hypothesis, we studied the distribution of both islands in the different MLST clonal groups of *A. baumannii*. The ST2 group was the one with the highest number of strains in our pangenome, as was to be expected due to its already known great evolutionary success (Fig. 4A). And particularly in this group, about two thirds of the genomes have the prophage DgiS1. On the other hand, the CRISPR-Cas I-Fa system was more represented in the ST79 group, although this group also presents genomes with DgiS1 (Fig. 4B).

**Fig. 4.**
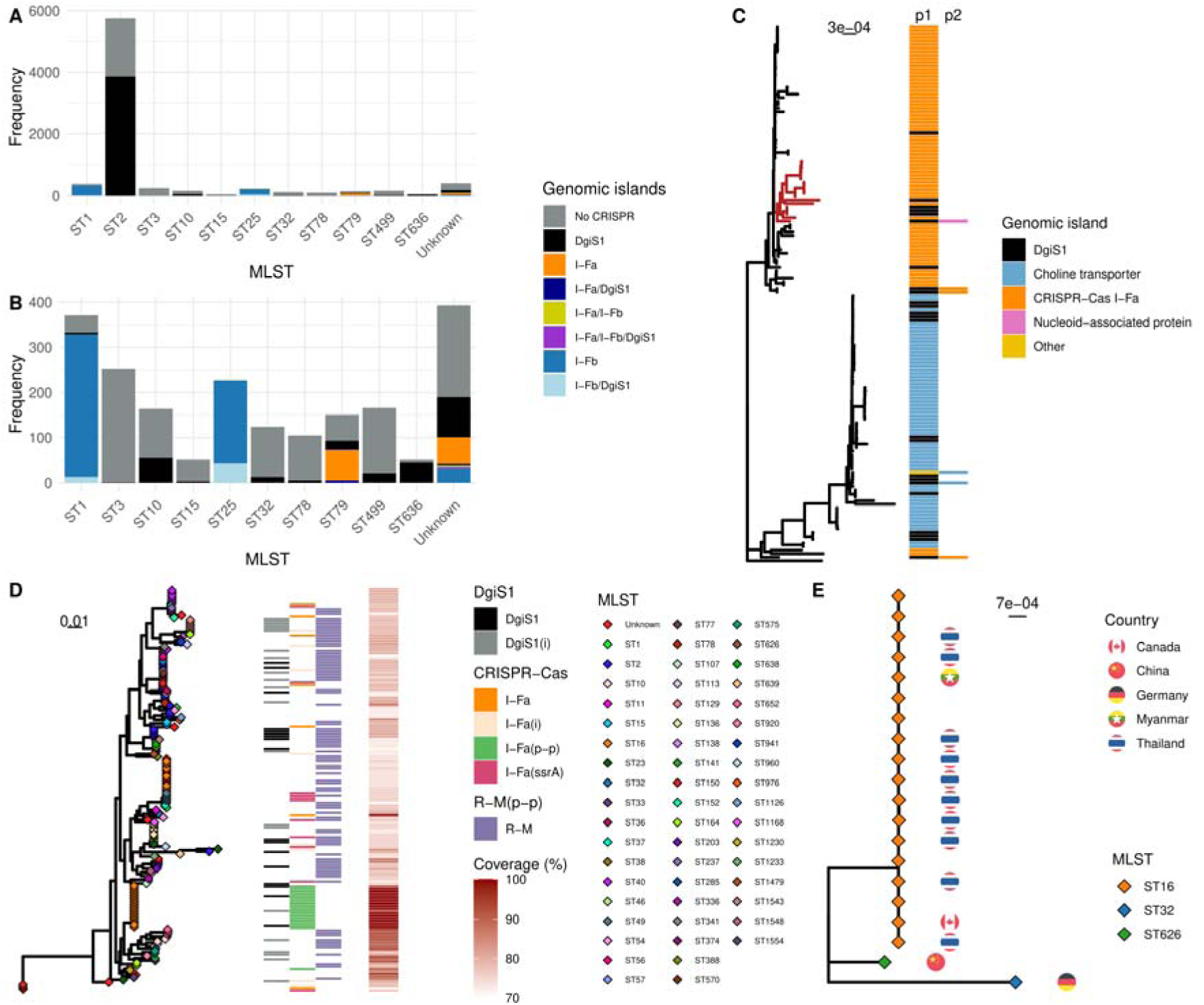
Distribution of genomes by MLST, and phylogeny of ST79 and the phage-plasmid PPTOP. (A) Distribution of genomic islands along the most abundant MLST groups of *A. baumannii*. “Unknown” corresponds to genomes to which it has not been possible to assign an MLST group. (B) Same figure as (A) but without showing ST2, to highlight the rest of the groups. (C) Molecular phylogeny of all genomes of the ST79 group, using as outgroup the reference genome of *A. baumannii*, which belongs to the ST46 group (the distance of this outgroup has been divided by 10, for better visualization). Column p1 shows the first integrated genomic island next to the *ssrA* gene, and column p2 the second integrated one. Behind the genomic islands shown is usually the core lipoprotein gene. A clade containing genomes with the DgiS1 prophage or the CRISPR-Cas I-Fa system on p1 is highlighted in red, whose root genome contains the Nucleoid-associated protein genomic island in the second position. (D) Molecular phylogeny of all phage-plasmid sequences found, from 70% sequence coverage with respect to a complete one. It is shown: the MLST groups in which the phage has been found, whether the genome carrying PPTOP also has DgiS1 (i=incomplete), or a CRISPR-Cas I-F system (i=incomplete, only a CRISPR array or a *cas* gene has been found; p-p=the CRISPR-Cas system is located in the phage-plasmid PPTOP; ssra=the CRISPR-Cas system is located next to the gene *ssrA*), or a restriction-methylation system (R-M). It is also shown the coverage of the phage-plasmid sequence that was found. (E) Molecular phylogeny of all phage-plasmids PPTOP found that carry the CRISPR-Cas I-Fa system in their sequence, included RefSeq:CP034093.1. The MLST groups in which the phage-plasmid has been found and the country of isolation are shown.

Thus, we took the ST79 group as the one that most frequently presented genomes with one or the other of the two genomic islands. The 152 genomes of this group are separated into two major clades (Fig. 4C). One of them contains genomes in which the CRISPR-Cas I-Fa system is very frequently integrated next to the *ssrA* gene. In the other clade, the genomic island with the choline transporter predominates in this first position. In contrast, the DgiS1 prophage appears 24 times distributed between both clades.

Comparing the genomes of the DgiS1-containing group with those containing the CRISPR-Cas I-Fa system, we could see that the main difference involved the genes of the opposite genomic island. Mostly, the presence of one of the islands seems to exclude the other. In only 3 genomes do both elements appear together, always with DgiS1 in the first position and the CRISPR-Cas system in the second. But it should be noted that the CRISPR spacers in these genomes do not target the DgiS1 sequence. All this would support the incompatibility of both genomic islands, even in very close genomes, and suggests that the DgiS1 prophage can sometimes escape the CRISPR-Cas system, displacing it downstream.

### The phage-plasmid PPTOP has an alternative defense system and appears in diverse genomes

On the other hand, we also wanted to know in what type of genomes the phage-plasmid PPTOP, which carries the CRISPR-Cas I-Fa system, was present. For this purpose, we searched for the PPTOP sequence, considering a threshold of 70% sequence coverage (at least 77 Kb). The sequence of this phage-plasmid was found in 164 genomes, again in the form of independent episomes. It appeared in several different clonal groups and seems to have been isolated in different countries, with or without the defense system (Fig. 4DE). Within these genomes, those carrying the CRISPR-Cas system usually do not have the DgiS1 prophage. Only six genomes had both elements, but in 4 of them the CRISPR-Cas system was present in the phage-plasmid PPTOP (not integrated with the *ssrA* gene), and in the others the system was incomplete (some or all of the *cas* genes were missing).

Interestingly, most of the remaining phage-plasmids PPTOP carry a type I restriction-methylation system that appears to alternate with the CRISPR-Cas I-Fa system. In addition, two of these PPTOP variants present the same att sequence that appears to use DgiS1. This att site was found next to the restriction-methylation system, suggesting that this new found defense system could also become integrated into the *ssrA* gene. This makes us propose this tmRNA gene as a hotspot for sequence exchanges and horizontal transfer, highlighting its role in defense.

### The CRISPR-Cas I-Fa systems of *A. baumannii* show common features regardless of their site of integration

To analyze the possible differences between the two CRISPR-Cas I-F subtypes, we counted the number of spacers in each of them. In the case of I-Fa, since it had two CRISPR arrays, these were analyzed separately. We considered CRISPR 1 array (C1a) the one closest to the *cas1* gene and CRISPR 2 array (C2a) the one closest to the *cas2_3* gene. Thus, the CRISPR arrays with the highest average number of spacers were C1a and those of the CRISPR-Cas I-Fb system (Fig. 5A). Within the CRISPR-Cas I-Fa system, C1a has almost twice as many spacers on average as C2a, something that holds true for the I-Fa systems associated with the choline transporter genomic island and the transposon. The trend was the opposite in CRISPR arrays carried by the phage-plasmid PPTOP. This could suggest that the CRISPR array grows, acquiring more spacers, when the CRISPR-Cas system is integrated into the bacterial chromosome, especially in the C1a. However, the low number of spacers appearing in PPTOP would be expected due to the space limitations the virus would have in its capsid.

**Fig. 5.**
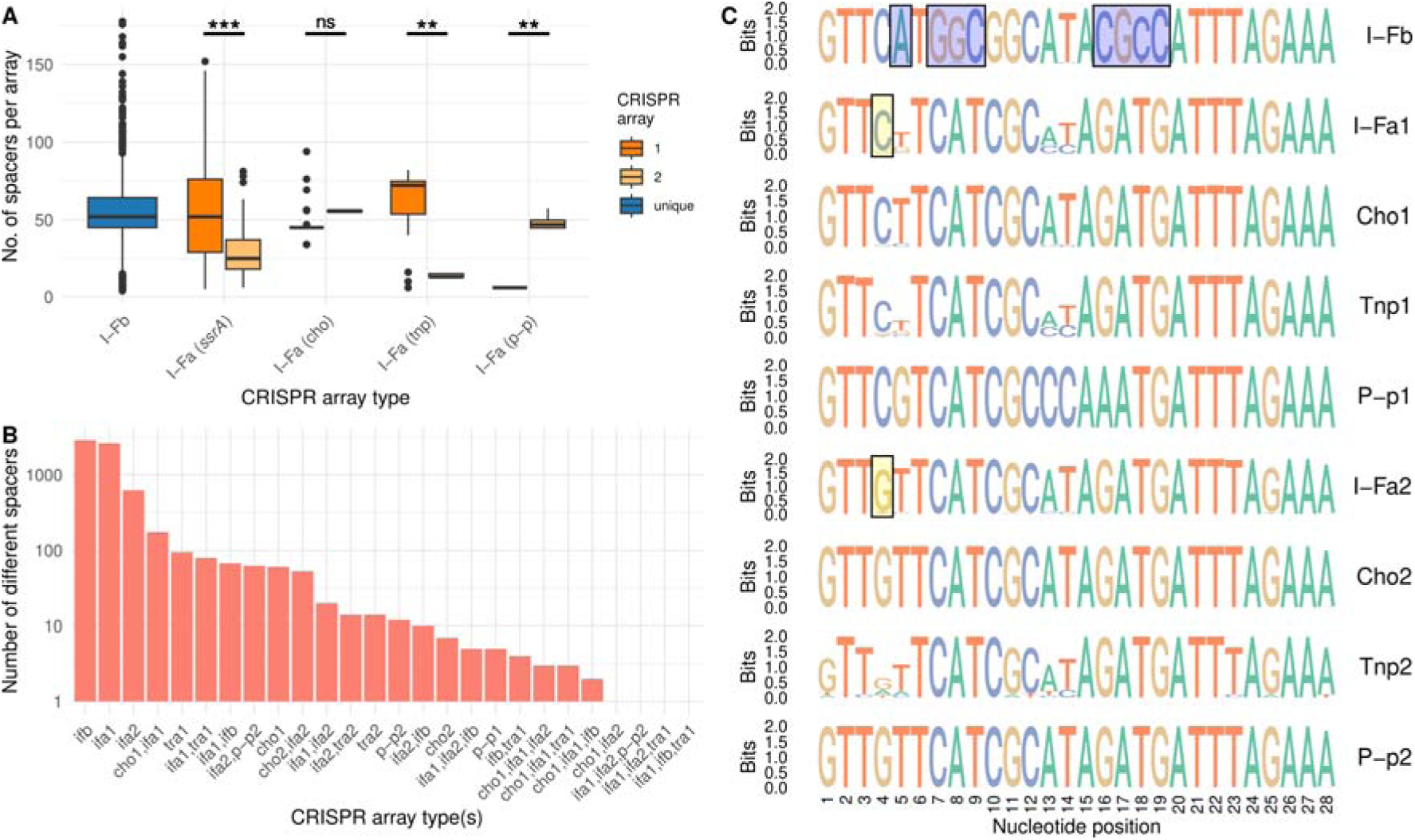
Number and type of spacers, and consensus sequence of repeats. (A) Number of spacers per CRISPR array separated by CRISPR-Cas system type; ifa=CRISPR-Cas I-Fa, ifb=CRISPR-Cas I-Fb, cho=choline transporter near to the ssrA, tnp=transposon near to the CRISPR-Cas I-Fa, p-p=CRISPR-Cas system in the phage-plasmid, 1=C1a near to *cas1*, 2=C2a near to *ca3_2*, unique=array of I-Fb. Asterisks highlight pairs of CRISPR arrays with significant differences in the median according to a Wilcoxon rank-sum test (from *≤0.05 to ***≤1e-16). (B) Number of non-redundant spacers separated according to the type of CRISPR-Cas system to which the CRISPR array belongs. When the same spacer appears in different types, they are separated by comma. The y-axis is in logarithmic scale. (C) Logo of the sequence repeats of CRISPR arrays separated by CRISPR-Cas system type. The differential I-Fb nucleotides have been shaded with blue boxes, and those distinguishing the two I-Fa arrays in yellow.

The number of different spacers is highest in the CRISPR arrays of the I-Fb system, followed closely by C1a (Fig. 5B). On the other hand, C2a has almost 5 times fewer than the other two. As expected, some spacers are common among the different systems, especially between C1a or C2a of the different I-Fa variants.

Finally, the nucleotide sequences of the repeats of the CRISPR-Cas systems were analyzed, and it was found that the I-Fb system differs from the I-Fa in 8 of its 28 nucleotides (Fig. 5C). However, the repeats of the two I-Fa systems differed by only 1 nucleotide (2 nucleotides in the case of C1a in PPTOP). This fact supports the idea that the C1a of all the different configurations of the CRISPR-Cas I-Fa system would have a common origin, as would all the C2a, on the other hand. Based on these results, the different CRISPR C1a CRISPRs will be considered as the same type of CRISPR-Cas I-Fa1 array regardless of its location from now on, and the same with the different C2a CRISPR.

### CRISPR-Cas I-Fa system spacers target all variants of the prophage DgiS1

The prophage DgiS1 is integrated into the *ssrA* gene and is present in a complete form in 2,199 genomes, with an average length of 79,989±7,846 bp. All of them can be considered as variants of the prophage. Moreover, the genomes analyzed have other prophages, which are also found in the form of different variants. To compare the interaction of the CRISPR-Cas systems with DgiS1 and with the rest of the prophages found in the bacteria, all the prophages found in the *A. baumannii* genomes were obtained and grouped by sequence similarity, finding a total of 330 different prophages. Of these, only the 40 that appeared in at least 100 genomes were taken for further analysis.

As expected, when analyzing the genomes in which such prophages appeared, including the DgiS1 prophages, it can be seen that these tend to be those lacking CRISPR-Cas systems (Suppl. Fig. S2). In fact, 18% of the genomes with a CRISPR-Cas I-Fa system, and 11% of those with the I-Fb type did not have any of these prophages. Whereas this was only the case in the 5% of genomes lacking CRISPR-Cas systems.

We then analyzed which prophage sequences were targeted by the spacers of the 3 different types of CRISPR arrays. The first thing that was found is that all DgiS1 variants were recognized by the I-Fa C1a spacers, and almost all variants (96%) by the C2a spacers (Fig. 6; Suppl. Fig. S3). However, hardly any DgiS1 variants are targeted by spacers of the CRISPR-Cas I-Fb system (6%). In addition, almost 85% of C1a genomes contain spacers against the prophage DgiS1, as do 67% of C2a genomes. And these spacers account for 4-5% of the different spacers in these CRISPR arrays. The second thing to note is that while the I-Fb spacers seem to recognize many of the different prophages (31 out of 41, if we consider 100% of their variants), these spacers do not target hardly any variants of two prophages: the referred DgiS1 and the Phage_1022.

**Fig. 6.**
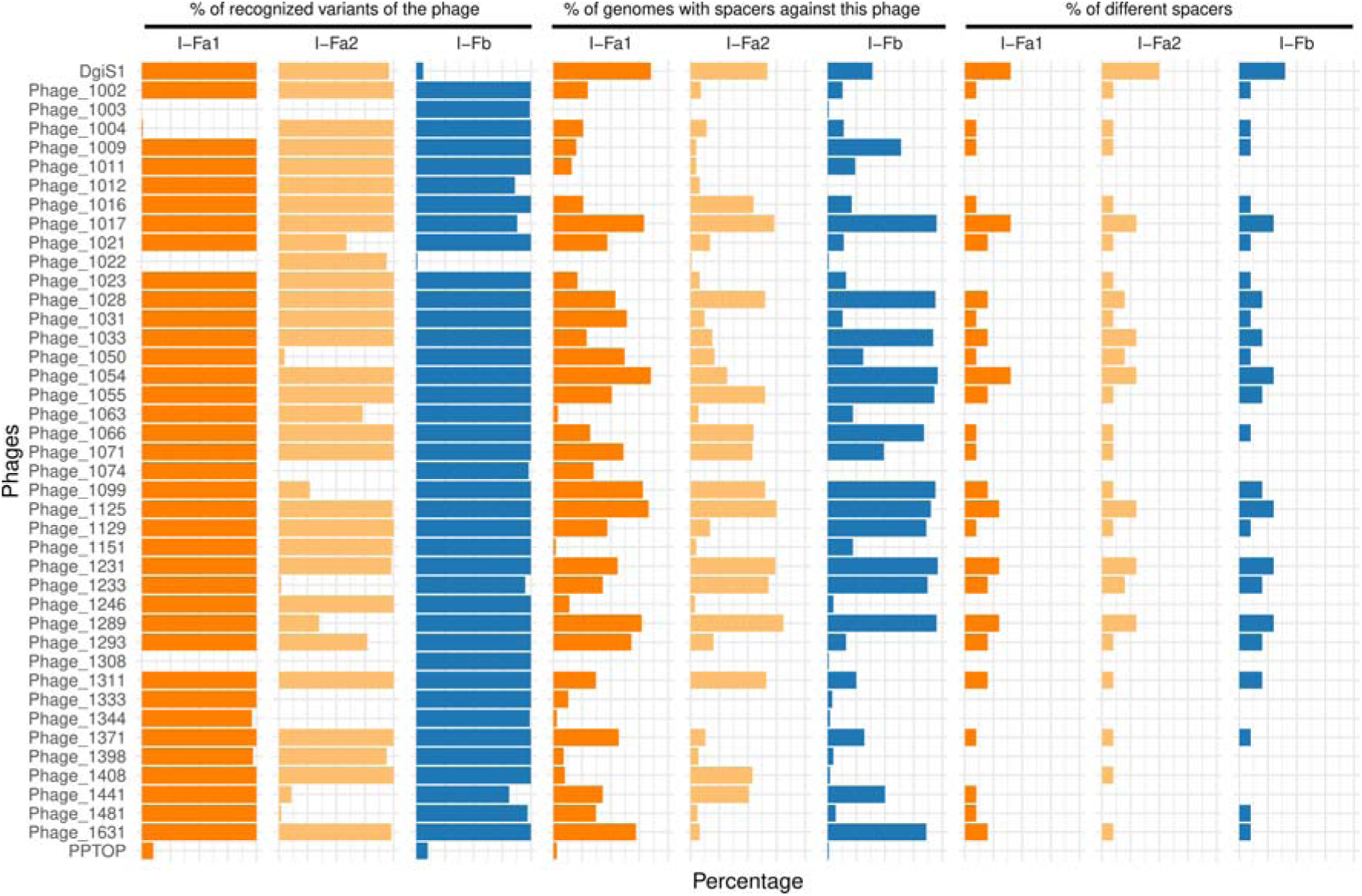
Spacers versus phages separated by CRISPR array type. The first block of 3 columns shows the proportion of different variants of each phage that are targeted by spacers of each type. Those in the second block show the proportion of bacterial genomes that have spacers against each of the phages. And those in the third block show the percentage of non-redundant spacers of each CRISPR type with respect to the total number of different spacers of that type. In all cases the X-axis shows the ratio between 0-100%, except in the third block, where it is shown between 0-10%, due to its lower values.

Notably, 44% of the CRISPR-Cas I-Fa array spacers target all 41 of these phages. However, only 18% of the spacers of CRISPR-Cas I-Fb systems target these prophages. This suggests some specialization on frequent phages by the I-Fa system. On the other hand, the spacers of the phage-plasmid PPTOP target 17 prophages, among which DgiS1 is not found. But this could be due to the low variability of the CRISPR arrays in the PPTOP variants that we have (5 different spacers in C1a and 76 in C2a). Although another explanation could be that the spacers against this virus were only acquired when the CRISPR-Cas system is integrated into the bacterial chromosome.

Finally, the variants of the phage-plasmid PPTOP were rarely targeted by any of the CRISPR-Cas systems. However, some variants were recognized by a few C1a spacers. The main spacer that targeted PPTOP came precisely from C1a of the phage-plasmid itself. This spacer matches 100% identity with the coding region of the *cas2_3* gene of the CRISPR-Cas I-Fa system. In fact, it highlights the idea that the homologous region of the gene *cas2_3* from the CRISPR-Cas I-Fb system is also the most conserved region of that gene, and in both *cas2_3* genes, the protospacer has the PAM sequence conserved (CC) almost 100% (Suppl. Fig. S4).

### The prophage DgiS1 could escape the CRISPR-Cas I-Fa system by mutations in its PAM sequences

The fact that multiple spacers were found against multiple sequence variants of the prophage DgiS1 suggests that this virus could be escaping the CRISPR-Cas system by fixing mutations in the protospacer or the corresponding PAM sequence. This would in turn force the bacterium to generate new spacers against the same virus.

To test this, we searched for the protospacers targeted by the spacers of the different types of CRISPR arrays. First, we searched for those that targeted sequences of the prophage DIGIS1. In the case of the CRISPR-Cas I-Fb system very few protospacers were found (454 protospacers), as had been seen before, but most of them conserved the expected PAM sequence CC (Fig. 7A). In the case of C2a more protospacers were found (2,208 protospacers), but most targeted the same sequence, again retaining the PAM sequence with the CC consensus. Finally, in the case of C1a the number of spacers increased almost 7-fold with respect to C2a (15,800 protospacers), and although the main PAM sequence was still CC, it was no longer as conserved and both positions could change to A or T (Fig. 7B). Specifically, 25% of the PAM sequences do not retain the CC dinucleotide (4,006 protospacers). This suggests that the acquisition of mutations in the phage PAM sequence is favored to evade a cognate CRISPR-Cas system that seems to be specialized against it. This would also be supported by the number of protospacers present in the prophage DgiS1, which could also suggest that these are mutations to escape the bacterial defense system.

**Fig. 7.**
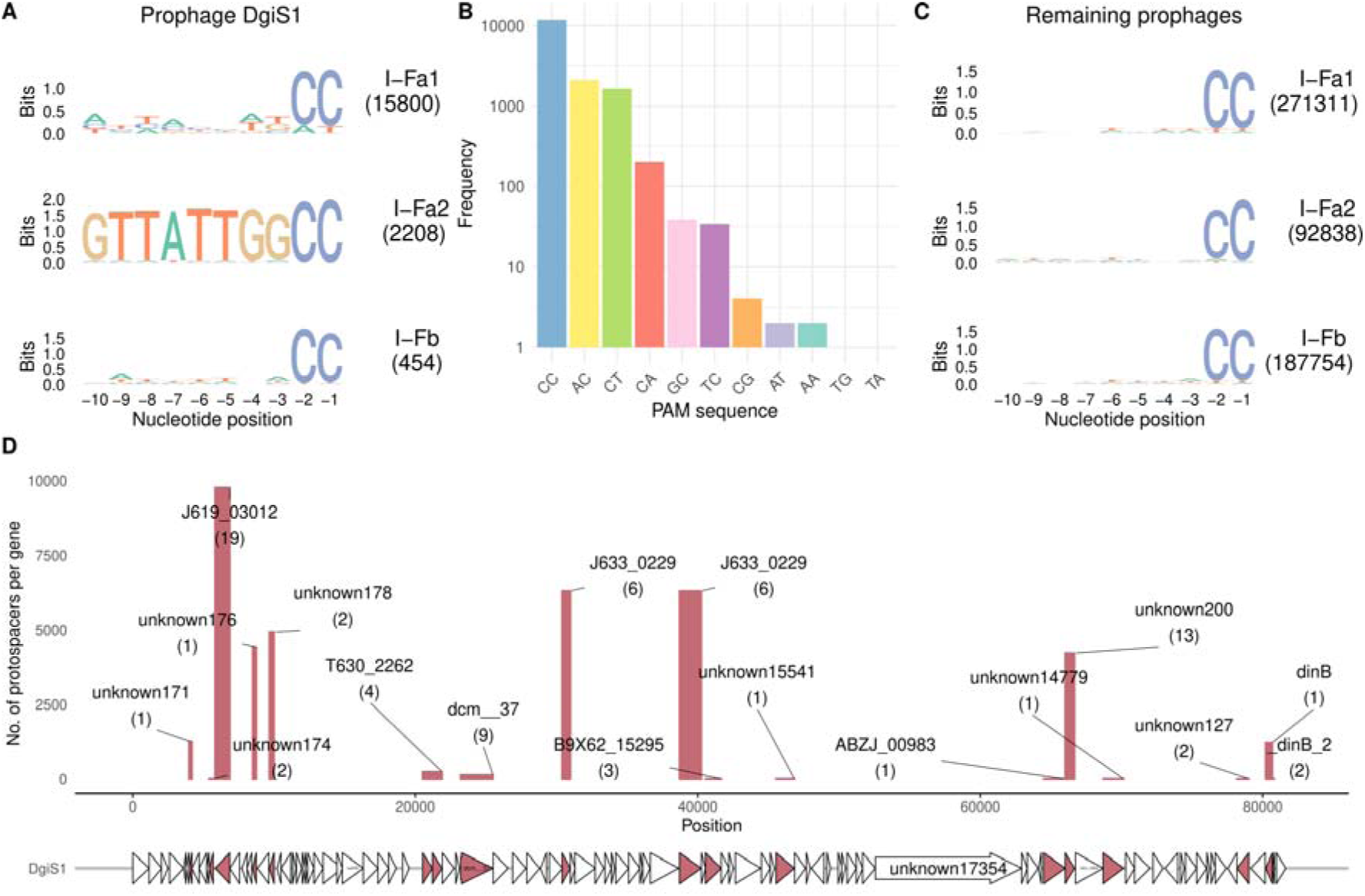
Spacers that recognize bacterial phages. (A) PAM sequence (10 nucleotides) upstream of the protospacers of the prophage DgiS1. The number in parentheses shows the number of different protospacers found. (B) Frequencies of consensus PAM sequences in the protospacers of the prophage DgiS1. The y-axis is in logarithmic scale. (C) The same as in (A) but with the rest of the bacterial prophages. (D) Number of protospacers found in the different genes of the prophage DgiS1. The number in parentheses below the gene name shows the number of protospacers that are different for that gene.

When the protospacers of the other bacterial prophages were analyzed, a high number of them were found in both CRISPR-Cas I-Fb and C1a, and approximately half in C2a (Fig. 7C). In addition, the PAM sequence was maintained as CC in most of the protospacers, with only 14% of them not conserving this dinucleotide. This result is significantly different from that found with the prophage DgiS1 (Fisher exact test, *p-value* = 1.10e-304).

Finally, to map the protospacers of the prophage DgiS1, all prophage genes appearing in the bacterial pangenome were taken and compared with the spacers of the CRISPR-Cas I-Fa system. Thus, protospacers could be found in 17 prophage genes, with several protospacers of different sequences appearing in most genes (Fig. 7D).

## Discussion

A small proportion of genomes of the pathogenic bacterium *A. baumannii* are known to exhibit two subtypes of type I-F CRISPR-Cas systems (8, 9). These were named as subtypes I-Fa and I-Fb. According to the most recent classification of CRISPR-Cas systems, the I-Fa subtype has been classified as I-F2, and the I-Fb subtype as I-F1. However, as we have found here, subtype I-Fa presents the *cas8f* gene, contrary to what has been described for subtype I-F2 (12). Nevertheless, these differences in the established configurations of these defense systems are commonly found in virus-borne CRISPR-Cas systems (18).

The knowledge of CRISPR-Cas systems in pathogenic bacteria, such as *A. baumannii*, is of interest. On the one hand, the presence of these systems can prevent the acquisition of resistance and virulence genes by the bacterium, having been found that genomes with I-Fa present a lower number of these genes than I-Fb, and even less than genomes without CRISPR-Cas systems (8, 15, 16). Although this relationship has not been confirmed in other studies (9). On the other hand, in the context of the currently flourishing phage therapy (use of phages to treat bacterial infections), it could be of special interest to know whether the strain that is producing the infection has effective defense systems against the phage to be used (31).

CRISPR-Cas systems are part of the accessory genome of prokaryotes, and it is known that they can be acquired or lost depending on the environment (8, 32). The manner in which these are acquired is not yet known. They have been associated with transposons and phages, which could serve as vectors to facilitate their horizontal transfer (33–35).

Here, we have found a CRISPR-Cas I-F system associated with a phage-plasmid (PPTOP), and this same defense system also appears to be integrated into a bacterial tmRNA gene. This integration site is a genomic archipelago in which different genomic islands appear integrated. Although some harbor alternative restriction-methylation defense systems, the site is mostly colonized by a prophage (DgiS1). When present, the prophage seems to exclude the CRISPR-Cas I-F system, and vice versa. In addition to this battle between the two elements for the same integration site, we have found that the phage-plasmid could be fighting against the prophage via the spacers of its CRISPR-Cas system, which appear to be specialized for DgiS1. But the latter would be defending itself, due to continuous mutations in its sequence, especially involving the PAM sequence of its putative protospacers. All this would demonstrate the warfare between the two phages and provide insight into how this CRISPR-Cas system spreads in *A. baumannii*.

In a previous study, in which this genomic archipelago was denoted as fGR4, the integration site was not found to be a tmRNA gene, but the prophage toxin-antitoxin system and the presence of methylases were found (36). And in another study, in which they named it as the island G13, they only mentioned the presence of a bacteriophage and methylation-restriction systems, in addition to other island genes (22). Nevertheless, this integration hotspot seems to be part of the *A. baumannii* defensome, being the battlefield of at least the two referred viruses.

The only phages characterized so far to contain a CRISPR-Cas system, which they use to defend itself against a second phage, are the *V. chloreae* phages ICP1 and the vibriophage 1.161 (19, 37). In the case of ICP1, both phages appear to compete with each other by also using the bacterium as a battlefield. The CRISPR-Cas system in ICP1 is also of type I-F, and presents a configuration similar to I-Fa of *A. baumannii* (except that *cas1* is rearranged), including the two CRISPR arrays. In addition, here we have found sequence similarity between their *cas* genes. However, so far it has not been reported that ICP1 integrated its CRISPR-Cas system into the bacterial genome, something that could be useful in the aforementioned phage warfare. In this case, the satellite virus is characterized as a phage-inducible chromosomal island (PICI). PICI are phage-derived satellite elements that integrate into the host chromosome, but are reactivated and excised upon phage infection and do not encode any capsid components, as they hijack those of the other phage (known as helper) (38). PICI are maintained in the bacteria, among other advantages because they prevent infection and lysis of the helper phage. The prophage DgiS1 found here could constitute a new PICI. In fact, some prediction tools only find an incomplete phage in its sequence. Although it should be noted that several genes putatively encoding capsid proteins are found in it.

We have found that the I-Fa system has a large number of spacers against the prophage DgiS1, especially from the array closest to the *cas1* gene (C1a). This array seems to acquire spacers against the different variants of the virus, which probably arise to escape from the defense system. The array could acquire the spacers against the prophage DgiS1 after the CRISPR-Cas I-Fa system is integrated into the bacterium. In addition, the array C1a appears to add new spacers against common phages in the bacterium, which could also positively impact the success of PPTOP by eliminating competitors.

Another way for phages to escape CRISPR-Cas systems, in addition to modifying their protospacer sequences, is by mutating the PAM sequence. This is something that has long been reported, and would be an efficient way to prevent recognition of the protospacer (39, 40). Specifically, phage ICP1 is less efficient in lysing *V. cholerae* when its satellite phage has non-canonical PAM sequences (30). Here we have seen that, although it may be a strategy common to all *A. baumannii* viruses targeted by CRISPR-Cas systems, it seems of particular relevance in the case of protospacers of the prophage DgiS1 targeted by the spacers of the C1a arrays. In addition, the presence of multiple spacers spanning different regions of the DgiS1 sequence may ensure virus neutralization by the CRISPR-Cas system, as has been demonstrated in other similar cases (30, 40, 41).

At least in the hospital environment, it seems that the phage-plasmid carrying the CRISPR-Cas system has not been able to gain the upper hand. ST2 strains of *A. baumannii* are the most commonly isolated in outbreaks of this bacterium, whereas ST79 and several other strains, which more frequently carry the CRISPR-Cas I-Fa system, are isolated more rarely, probably because their infections have a better prognosis (6, 42). This could be in agreement with the already described weaknesses that carrying such a large sequence as a CRISPR-Cas system by a virus may entail, thus requiring alternative systems (43).

However, the CRISPR-Cas I-Fa system, and more specifically thanks again to its array C1a, seems to be an efficient defense against the most frequent phages that integrate into bacteria, apart from prophages DgiS1. This, indirectly, would also benefit the phage-plasmid that carries this defense system.

The CRISPR-Cas I-F system of the phage ICP1 suppresses the antiphage function of the satellite but the mechanism used for this is still under study (44). We have found that some phage-plasmids carrying the CRISPR-Cas I-Fa system carry spacers against the most conserved region of the coding sequence of the *cas2_3* gene, both of the I-Fa subtype itself and of the I-Fb subtype. This could be a clue to their function. These anti-cas spacers have been seen previously in studies with other species, but their function has not been determined (45). Although its high conservation suggests that such a specific function must exist.

This particular spacer appears in the array C1a of the phage-plasmid PPTOP, which has a smaller number of spacers. In addition, the C1a arrays have repeats that differ from the C2a arrays in the nucleotide in the fourth position. And it has already been shown that variations in the sequence of the repeats can lead the CRISPR system to act as a regulator of endogenous genes, including *cas* genes (46). This is common in so-called mini-arrays, composed of 2 repeats (sometimes with a degenerate sequence) and a single spacer, and usually appear in the vicinity of complete CRISPR-Cas systems. These mini-arrays are often found in phages and are often associated with blocking the superinfection of the phage itself. Likewise, anti-cas spacers located in orphan I-F CRISPR have been described in *Escherichia coli* as a system to prevent the acquisition of full CRISPR-Cas systems (47). The fact that these CRISPR arrays are short and appear close to the main CRISPR array could explain why some phages, such as ICP1 and PPTOP, have two CRISPR arrays with an unequal number of spacers.

In addition, PPTOP sometimes appears to carry alternative defense systems to CRISPR-Cas, such as a restriction-methylation system. Alternative systems to fight against its rival virus have also been described in the case of phage ICP1, but with a recombinase system involved (48, 49). This fact suggests that both viruses are adapting in a guns and hire race in which each one can impose variants, which are eventually overcome by the other virus and force both to continue adapting (41, 43).

In conclusion, it seems increasingly clear that anti-phage systems can be used by the phages themselves to eliminate competitors. In this case, these defense systems are carried by a phage-plasmid that although it would be targeted towards the most frequent viruses of the bacterium *A. baumannii*, it seems to be especially directed towards a very frequent prophage in the species. This frequency suggests that, at least in this specific warfare, carrying such weapons is not enough to succeed in the artificial hospital environment to which we have brought this pathogenic bacteria.

## Acknowledgements

We thank C3UPO for the HPC support, and Dr. Rubén de Dios Barranco for the critical reading of the manuscript.

## Data and Materials Availability

All data needed to evaluate the conclusions in the paper are present in the paper and/or the Supplementary Materials.

The code used to analyze the data is available in the repository: https://github.com/UPOBioinfo/cancun

