## Supplementary figures and images for "Biological warfare between two bacterial viruses in a genomic archipelago sheds light on the spread of CRISPR-Cas systems"

### Suppl. Fig. S1

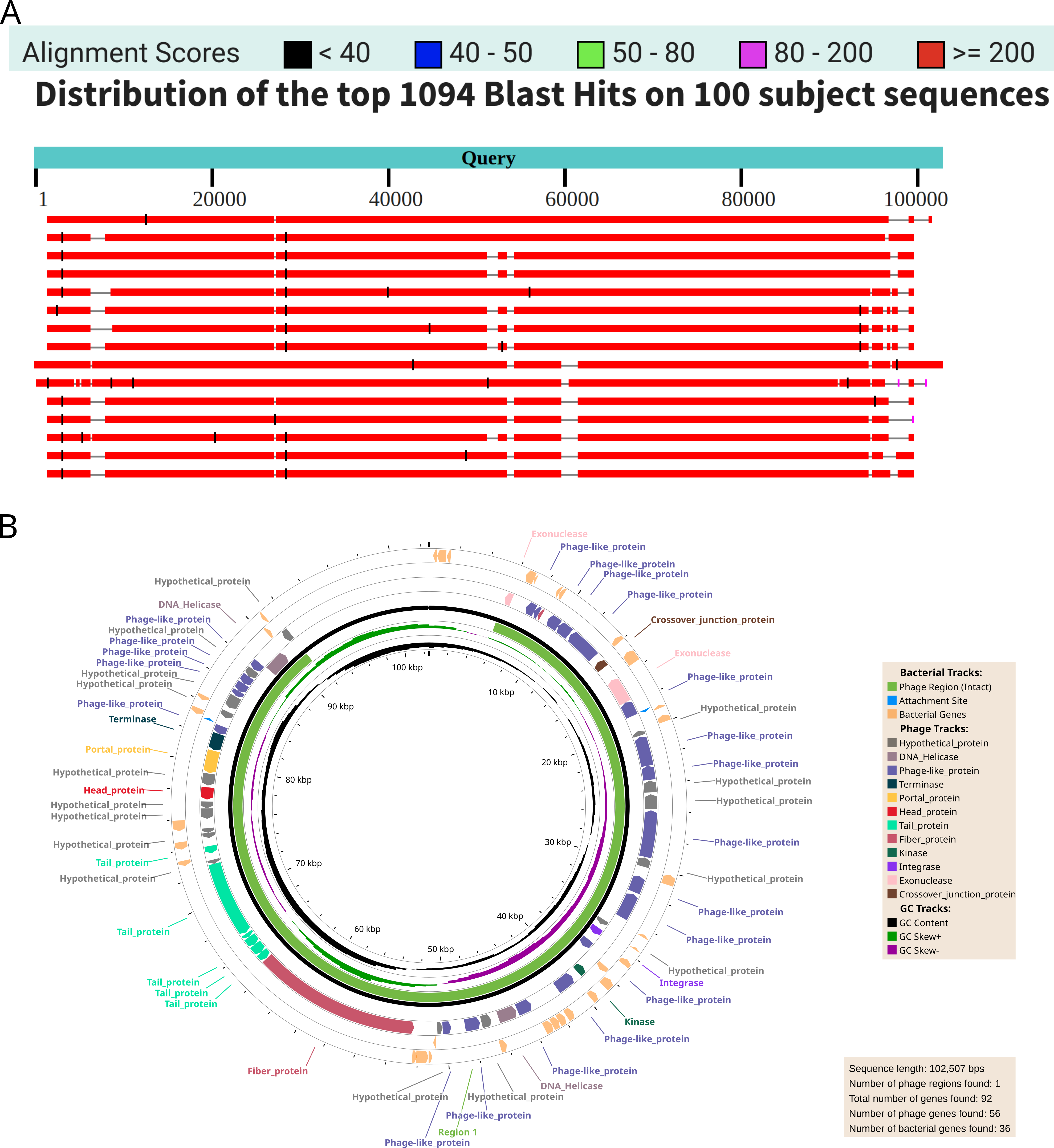

### Suppl. Fig. S2

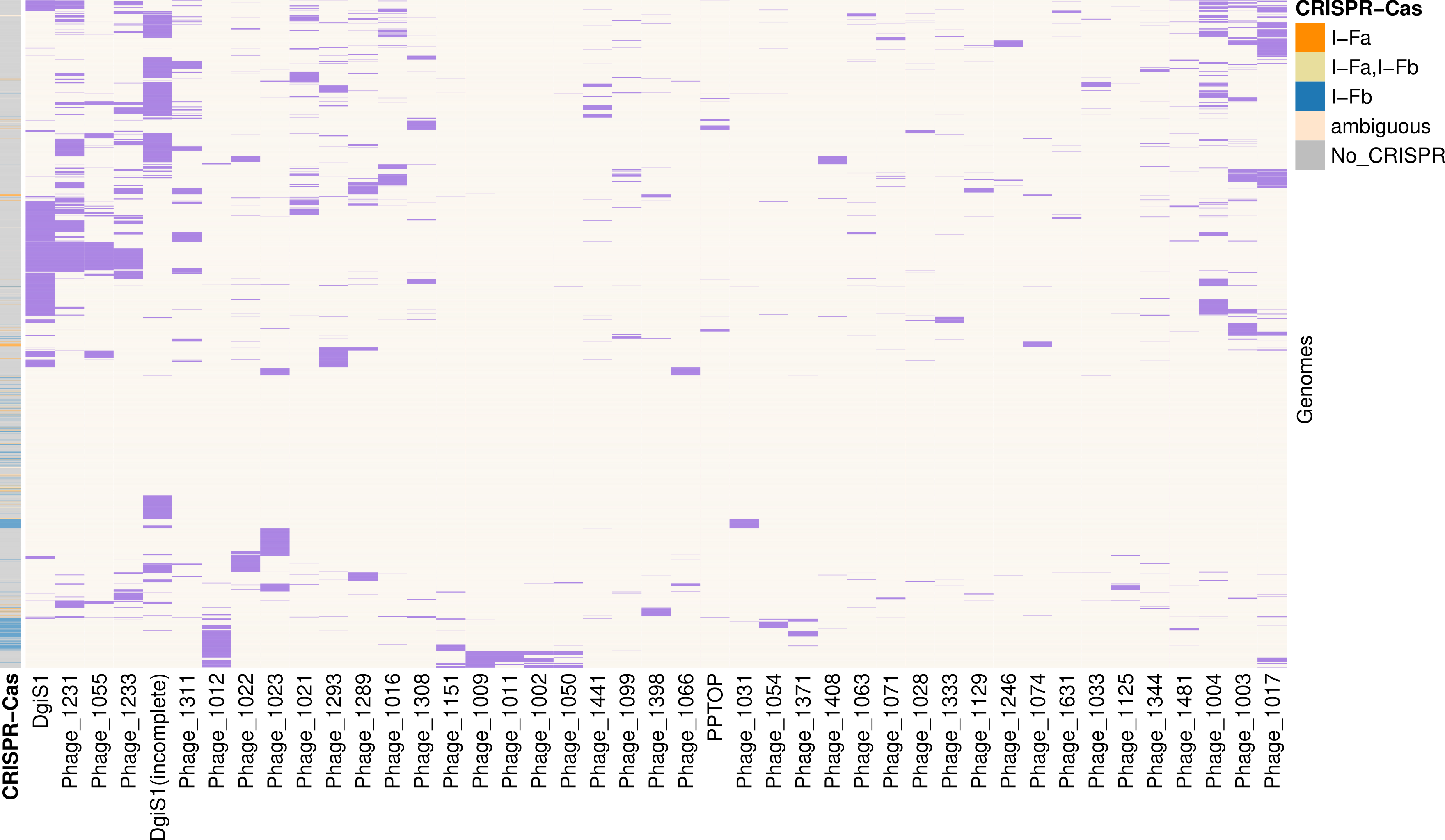

### Suppl. Fig. S3

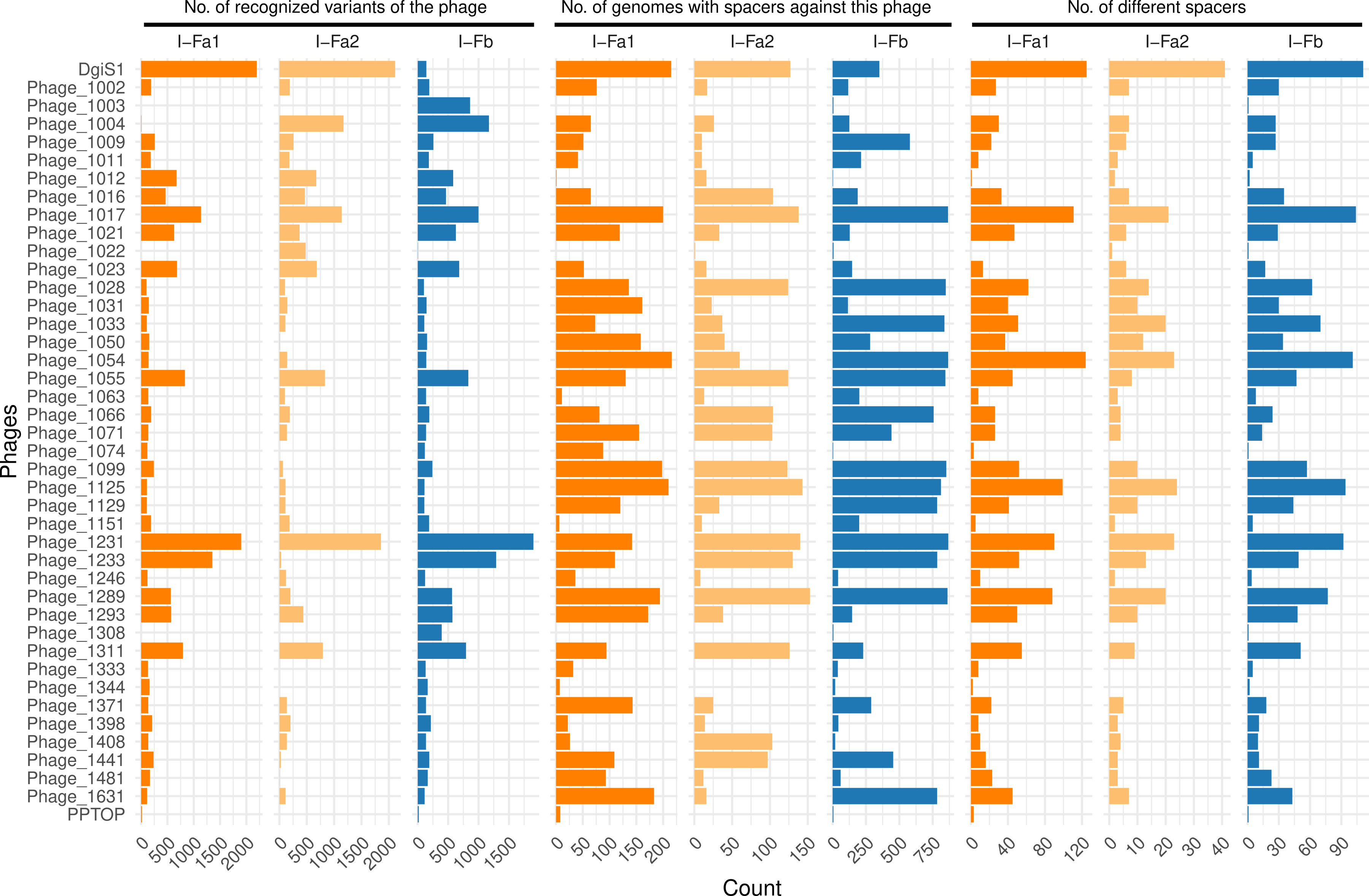

### Suppl. Fig. S4

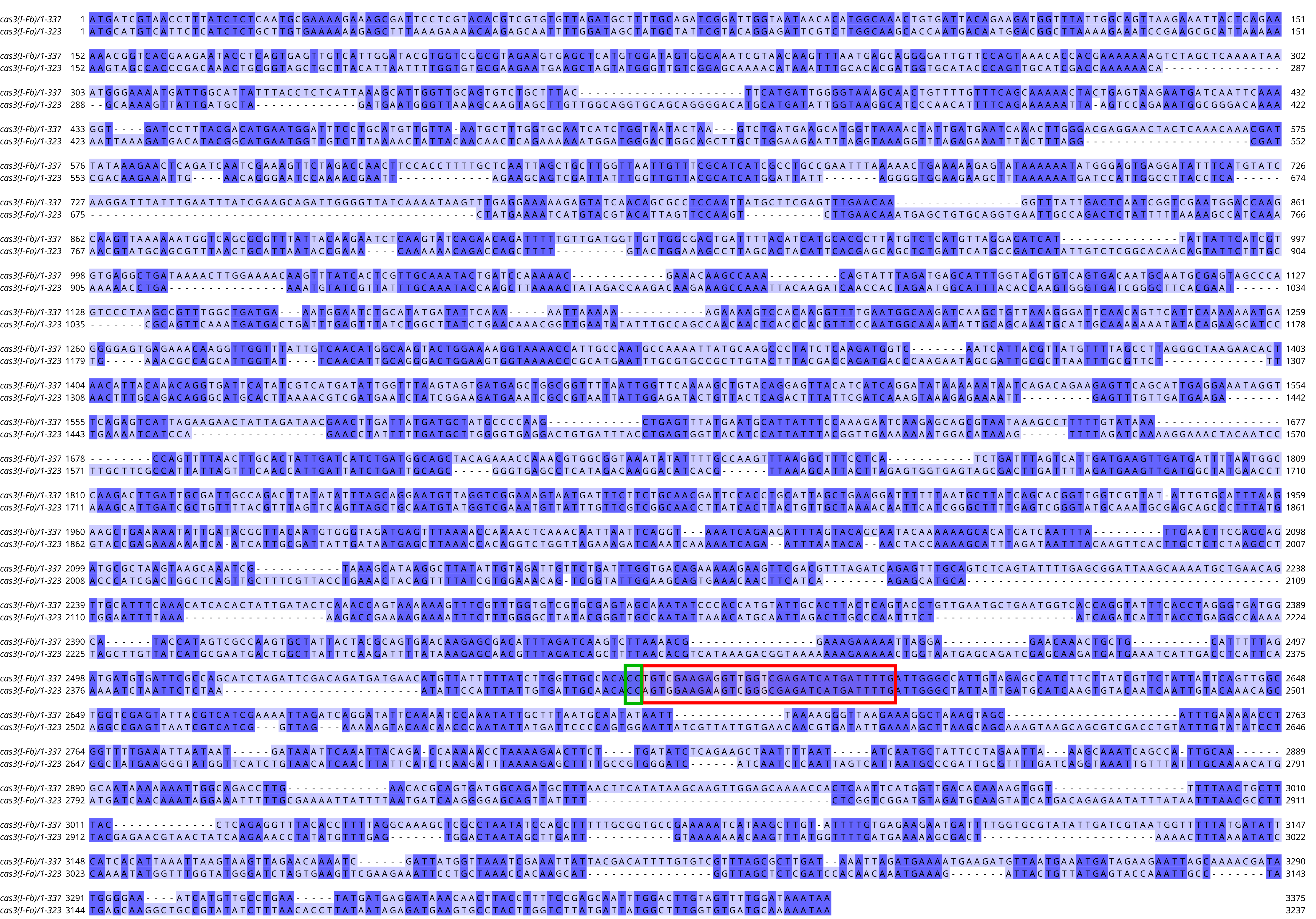
